# Precise Alternation Between Image-Forming Sample Planes Enables Quantitative Monitoring of Receptor-Arrestin Interaction Dynamics at the Plasma Membrane of Live Cells

**DOI:** 10.64898/2026.04.15.718721

**Authors:** Thomas D. Killeen, Michael R. Stoneman, Ionel Popa, Qiuyan Chen, Valerică Raicu

## Abstract

Investigations of G protein-coupled receptors (GPCRs) interactions with non-visual arrestins in living cells are essential to understanding the complex molecular mechanisms of GPCR-based signaling. Quantitative analysis of these interactions remains challenging in live cells, particularly when attempting to repeatedly image distinct cellular regions with high precision. Here, we describe the implementation of an optical imaging stabilization approach that integrates the recently developed Focal Readjustment for Enhanced Vertical Resolution (FREVR) technology into a multiphoton microscope, enabling high-precision alternation between image-forming sample planes with < 20-nanometer repeatability and stability over time. Using this setup, we monitored the dynamic recruitment of arrestin-2 (Arr2) to the plasma membrane of HEK-293 cells expressing muscarinic acetylcholine M_2_ receptors (M_2_R) by alternatively imaging distinct planes of interest, the basolateral membrane and a membrane cross-section. Following stimulation of M_2_R by agonist ligand, we observed a pronounced redistribution of cytoplasmic arrestin-2 toward the plasma membrane in both cellular cross-sections and at the basolateral membrane. This method enables direct comparison of receptor and arrestin dynamics across regions of individual cells with very high precision, eliminating the need for averaging over numerous cells in order to denoise biologically relevant signals, and thereby capturing physiological cell-to-cell variability.

## 1. INTRODUCTION

G protein-coupled receptors (GPCRs) mediate a vast array of cellular signaling processes, regulating physiological responses to hormones, neurotransmitters, and environmental stimuli^1–3^. GPCRs constitute one of the largest and most therapeutically relevant classes of membrane proteins^4, 5^. However, despite decades of study, our understanding of the complex mechanisms and diverse protein interactions involved in GPCR signaling remains incomplete. Among these interactions, the relationship between GPCRs and the ubiquitously expressed proteins known as non-visual arrestins (arrestin-2 and arrestin-3) has garnered significant attention^6, 7^. Beyond their established roles in receptor desensitization and internalization, arrestins are now recognized as multifunctional scaffolds that initiate distinct signaling pathways^8–10^. As research into these signaling functions advances, the need for quantitative tools capable of resolving the spatial and temporal dynamics of arrestin recruitment in living cells becomes more acute.

Among the most useful techniques for probing GPCR signaling in live cells are advanced fluorescence microscopy approaches, which offer the spatial and temporal resolution needed to resolve dynamic protein trafficking and interactions. Recent innovations in this field^11^, including Förster Resonance Energy Transfer (FRET) spectrometry^12–14^ and Fluorescence Intensity Fluctuation (FIF)-based analysis approaches^15–17^, have provided direct insight into protein association and receptor oligomerization in live cells. Fluorescence microscopy studies have further demonstrated that arrestin recruitment to the plasma membrane involves spatially regulated molecular interactions that require accurate tracking of protein redistribution across distinct cellular compartments^18, 19^.

Our previous investigation of the interaction between fluorescently tagged muscarinic acetylcholine M^2^ receptors (M_2_Rs) and non-visual arrestins at the basolateral plasma membrane of human embryonic kidney (HEK-293) cells^19^ demonstrated the feasibility of monitoring the dynamics of arrestins and receptor interactions and their co-localization at the plasma membrane, albeit with significant technical limitations, as follows. (1) Axial positioning uncertainty and the temporal drift of the basolateral membrane out of the microscope’s focal plane, caused by inherent mechanical and thermal instability in both the imaging system and the sample, introduced significant variability in those measurements. In order to reduce this inherent noise, it was necessary to average data from multiple cells to extract meaningful trends in receptor and arrestin behavior. However, because cells often differ in receptor and arrestin expression levels as well as in their physiological state—for example due to being in different cell cycle phases during imaging—averaging across populations can obscure the dynamics occurring within individual cells. (2) A second challenge revealed by our previous study was that fully monitoring the dynamics of arrestin—a cytoplasmic protein—as it relocates from cytoplasm to the plasma membrane requires one to also image cross-sections deeper within the cells. This, in turn, requires alternating imaging planes between basolateral and cross-sectional views during the duration of the entire physiological processes involved (that can last for an hour or sometimes longer). Unfortunately, repeated switching between image planes dramatically expands the list of instrumental errors to include mechanical hysteresis, backlash in the stage drive, and even hydrodynamic effects as the objective moves through immersion medium. These effects make it difficult to reliable return to the exact same focal plane^20^.

To overcome both of these challenges, in the present study we implemented and adapted a recently developed technology, Focal Readjustment for Enhanced Vertical Resolution (FREVR)^21^, in a spectrally resolved two-photon micro-spectroscope^14^ which was built around a commercially available microscope that utilizes a standard objective stage actuator. Despite relying on a conventional stage, our present implementation maintains better than 20 nanometers axial positioning resolution during image acquisition, demonstrating that nanometer-scale focal positioning can be achieved without specialized piezo positioning hardware. Unlike conventional focus-lock systems that hold a single plane in focus^22, 23^, FREVR enables repeatable positioning across a wide range of focal depths with high precision.

Using this stabilized imaging system, we conducted proof-of-concept quantitative single-cell imaging of arrestin-2 (Arr2) redistribution between cytoplasm and the plasma membrane as well as arrestin-2 and M_2_R localization and co-localization at the plasma membrane following receptor activation with agonist ligand. We quantified the temporal evolution of Arr2 fluorescence at the plasma membrane following receptor activation, accompanied by a corresponding change in cytoplasmic signal. Together, these changes reflect the recruitment of cytoplasmic Arr2 to activated M_2_R at the plasma membrane. These results highlight how accessible active-stabilization methods can enhance the precision of quantitative fluorescence microscopy, enabling detailed analysis of single-cell signaling dynamics and receptor-signaling partner interactions.

## 2. MATERIALS AND METHODS

### 2.1 Sample dish preparation

Glass-bottom dishes consisting of a 35 mm plastic chassis with a 14 mm #1.5 cover glass micro-well (Fisher Scientific) were functionalized with 50 µg ml^-1^ poly-D-lysine (Gibco) according to the manufacturer’s protocol. Amino-coated polystyrene beads (3.75 µm diameter, Spherotech) were diluted in DPBS and added to the microwell to allow adsorption onto the glass surface. Beads were added after functionalization of the cover glass with poly-D-lysine, and prior to the addition of the biological sample of interest (HEK-293 cells). Each well was seeded with approximately 40,000 such reference beads, resulting in an average of ∼5 reference beads per field of view (approximately 0.01 mM_2_).

### 2.2 Cell line generation and culturing

#### 2.2.1 Stable cell line generation

To generate a stable cell line expressing arrestin-2 (Arr2), HEK-293 cells (human embryonic kidney cells; ATCC, CRL-1573) were transfected, using Lipofectamine^TM^ 3000 reagent (Thermo Fisher Scientific) following the manufacturer’s protocol, with plasmids encoding mCitrine-tagged Arr2 (mCitrine-Arr2)^24^. In this construct, mCitrine was fused at the N-terminus of the Arr2 open reading frame^24^. The plasmid also contained a Geneticin (G418) resistance cassette, enabling antibiotic selection. After transfection, cells were cultured in standard DMEM (high glucose; Gibco) supplemented with 10% (v/v) fetal bovine serum (FBS), 100 U ml⁻¹ penicillin, and 100 µg ml⁻¹ streptomycin for two days without antibiotic. Selective pressure was then applied by switching to growth medium containing 500 µg ml⁻¹ Geneticin. Cells were propagated until antibiotic-resistant colonies formed. These colonies were selected, propagated, and screened for mCitrine-Arr2 expression through examination of fluorescence spectra using the fluorescence microscope described in Section 2.3. Clones with detectable mCitrine fluorescence were used as a stable cell line for all subsequent experiments. These cells are referred to hereafter as *Arr2 cells*.

#### 2.2.2 Sample preparation

Arr2 cells were cultured as monolayers in T-25 flasks (CellTreat) using DMEM (high glucose) supplemented with 10% (v/v) fetal bovine serum, 100 U ml^-1^ penicillin, 100 µg ml^-1^ streptomycin, and 500 µg ml^-1^ Geneticin (Gibco). Cells were maintained in a humidified incubator with 95% air and 5% CO^2^ at 37°C. Two days prior to imaging, cells were subcultured and seeded in functionalized 35 mm dishes with adsorbed reference beads (see Methods Section 2.1) at a density of 2.0×10^5^ cells per dish. The following day, the cells were transfected with plasmids encoding muscarinic acetylcholine M_2_ receptor (M_2_R)^25^ fused at its N-terminus to monomeric enhanced GFP (mEGFP-M_2_R)^26, 27^. Transfection was performed using Lipofectamine^TM^ 3000 reagent (Thermo Fisher Scientific) following the manufacturer’s protocol. After transfection, the cells were incubated in a humidified incubator with 95% air and 5% CO^2^ for another 24 hours. Prior to imaging, the transfected cells were removed from the incubator, washed with Dulbecco’s phosphate-buffered saline (DPBS; Gibco) and maintained in DPBS for imaging.

### 2.3 Imaging setup

#### 2.3.1 Two-photon fluorescence microscope

A custom-built two-photon micro-spectroscope^14^ was used to acquire all fluorescence images presented in this study. The instrument consists of a Zeiss Axio Observer inverted microscope stand equipped with a C-Apochromat 63× water immersion objective (NA = 1.2; Carl Zeiss Microscopy). A femtosecond-pulsed mode-locked Ti:Sapphire laser (MaiTai^TM^, Spectra-Physics), was tuned to a wavelength of 960 nm, expanded using a Galilean beam expander, and reflected off a phase-only spatial light modulator (SLM; P1920-1152-HDMI Nematic, Meadowlark Optics) for excitation beam shaping. A phase-mask was imposed onto the SLM to generate a 6 x 4 array of staggered beamlets, similar to methods detailed previously^14, 28^, resulting in 24 voxels of excitation with an average power of 3.1 mW per voxel. The excitation array was scanned across the sample plane with a pixel dwell time of 5 ms using an in-line pair of galvanometric mirrors (galvos). The fluorescence emitted from the sample was projected through an optical transmission grating onto a cooled electron-multiplying charge-coupled device (EMCCD) camera (iXon Ultra 897, Andor Technologies) resulting in fluorescence spectra spanning 450 to 600 nm across multiple pixels. Custom software, written in C++, was used to control the Axio Observer, the SLM, the galvos, and the EMCCD. Data acquired from the EMCCD was reconstructed into a stack of images consisting of 188 x 128 pixels and 16 wavelength channels.

#### 2.3.2 Focal Readjustment for Enhanced Vertical Resolution

FREVR was implemented by inserting a notch laser dichroic beamsplitter (Semrock, NFD01-633-25×36), referred to as DM1 in Figure 1, into the Zeiss Axio Observer reflector turret. The FREVR module accessed the rear port of the microscope and included a dichroic mirror (DM2), a light emitting diode (LED), a Complementary-Metal-Oxide-Semiconductor (CMOS) camera, and optical filters that narrow the LED wavelength band. This implementation provides a unique focal plane for FREVR to operate within, decoupled from the focal plane of the EMCCD camera paired with the fluorescence microscope.

**Figure 1.**
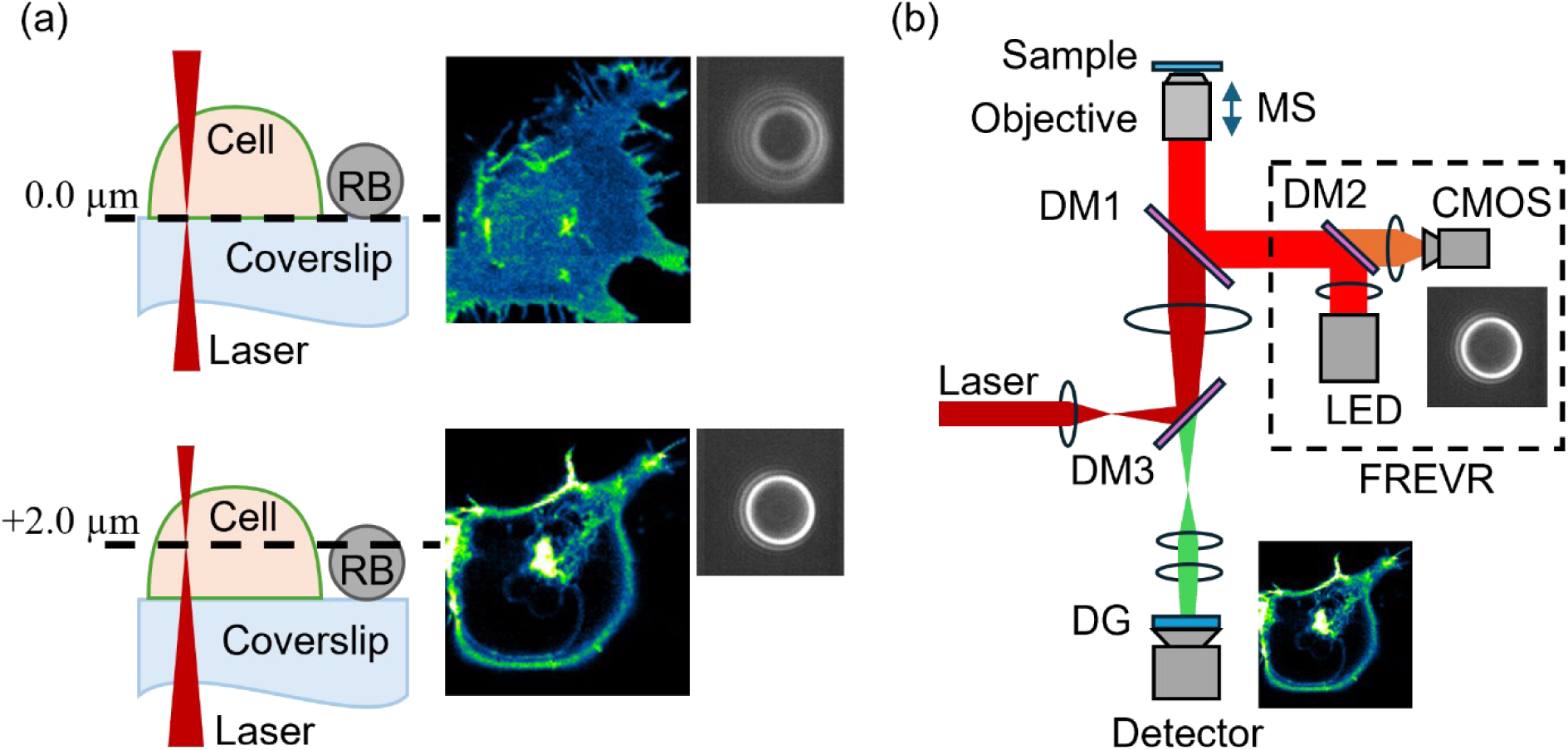
(a) Schematic illustration of a cell expressing fluorescently tagged proteins, imaged using FREVR at the basolateral membrane and a plane 2.34 µm into the cell, with representative fluorescence images and corresponding reference bead (RB) images shown. (b) Diagram of the inverted fluorescence microscope equipped with the FREVR module incorporating: Dichroic Mirrors (DM#), Motorized Stage (MS) Diffraction Grating (DG), Detector (EMCCD camera), light emitting diode (LED), and Complementary-Metal-Oxide-Semiconductor (CMOS) camera.

The objective stage of the Axio Observer was operated via custom control software implemented using the Zeiss SDK, allowing synchronized z-positioning and image collection. FREVR software^21^ was used to generate reference bead libraries, determine the bead position in the stack library, and apply focal corrections in real time. Custom functions were added to the C++ code to enable mechanical control of objective focus through the Zeiss SDK interface.

### 2.4 Sample imaging

#### 2.4.1 Imaging beads for axial positioning calibration

Z-positioning stage calibration was performed on the Zeiss Axio Observer microscope using 3.7 µm polystyrene beads (Spherotech: PP-35-10) and 2.8 µm magnetic beads (Dynabeads: 65305) immobilized on glass coverslips, together with glass surface impurities as fiducial markers. A 3D reflected-light image stack consisting of 500 frames from a single field of view was acquired by stepping the objective stage actuator in 20 nm increments and recording one frame after each step. For each bead, a central region of interest was selected, and horizontal and vertical intensity profiles were extracted through the bead center in every frame (Figures S1 and S2). A symmetric double-Gaussian fit was applied to each intensity profile to determine the bead width (in pixels) as a function of the reported objective position.

An axial calibration factor was then defined as the ratio between the true focal-plane displacement and the microscope-reported objective displacement. The axial calibration factor was determined using two independent approaches, as described in Supplementary Note 1. In the first method, the reported objective displacement between the glass surface and the center of each bead was measured and compared with the manufacturer-reported bead radius. To determine the glass surface position, we located a surface impurity adjacent to the bead, and a small region of interest was defined around it; a two-dimensional Gaussian fit was performed on each image frame in the z-stack (Figure S1). The reported objective position at which the fitted amplitude reached its maximum was taken to correspond to the glass surface (since the impurity was assumed to be sub-diffraction sized). In the second method, the reported objective displacement between the planes of the centers of the two differently sized beads in the same field of view was measured and compared with the known difference in bead radii. In both cases, manufacturer-reported bead dimensions were used to determine the correction factor relating reported stage position to true focal-plane displacement (Supplementary Note 1; Figures S1 and S2). These measurements yielded a consistent calibration factor of 1.17 across both measurement methods. All Zeiss objective position readouts were corrected accordingly and reported using this corrected scale throughout the remainder of the manuscript.

#### 2.4.2 Benchmarking the repeatability of axial positioning of the sample

Sample chambers containing fiducial reference beads, prepared as detailed in Methods Section 2.1, were imaged on the FREVR equipped microscope described in the previous section to quantify the precision of FREVR-assisted axial positioning relative to the motorized objective stage position readout of the Zeiss Axio Observer. A reference bead (RB) was located in the field of view, and a reference bead library was generated by imaging the RB at various focal planes, spanning 4.68 µm axial range with 23 nm spacing between reference bead images. The system was programmed to perform a mock imaging protocol which consisted of five sequential z-scans. The z-scans consisted of three planes of interest, a baseline plane, and two additional planes at 0.59 µm and 2.34 µm above the baseline. There was a programmed delay of 50 seconds between each z-scan. The measured focal plane position determined by FREVR, the motorized stage offset reported by the Zeiss Axio Observer, and the difference between the measured focal position and the target position commanded by the control program were recorded during acquisition to quantify the stability and uncertainty of the system (Figure 2).

**Figure 2.**
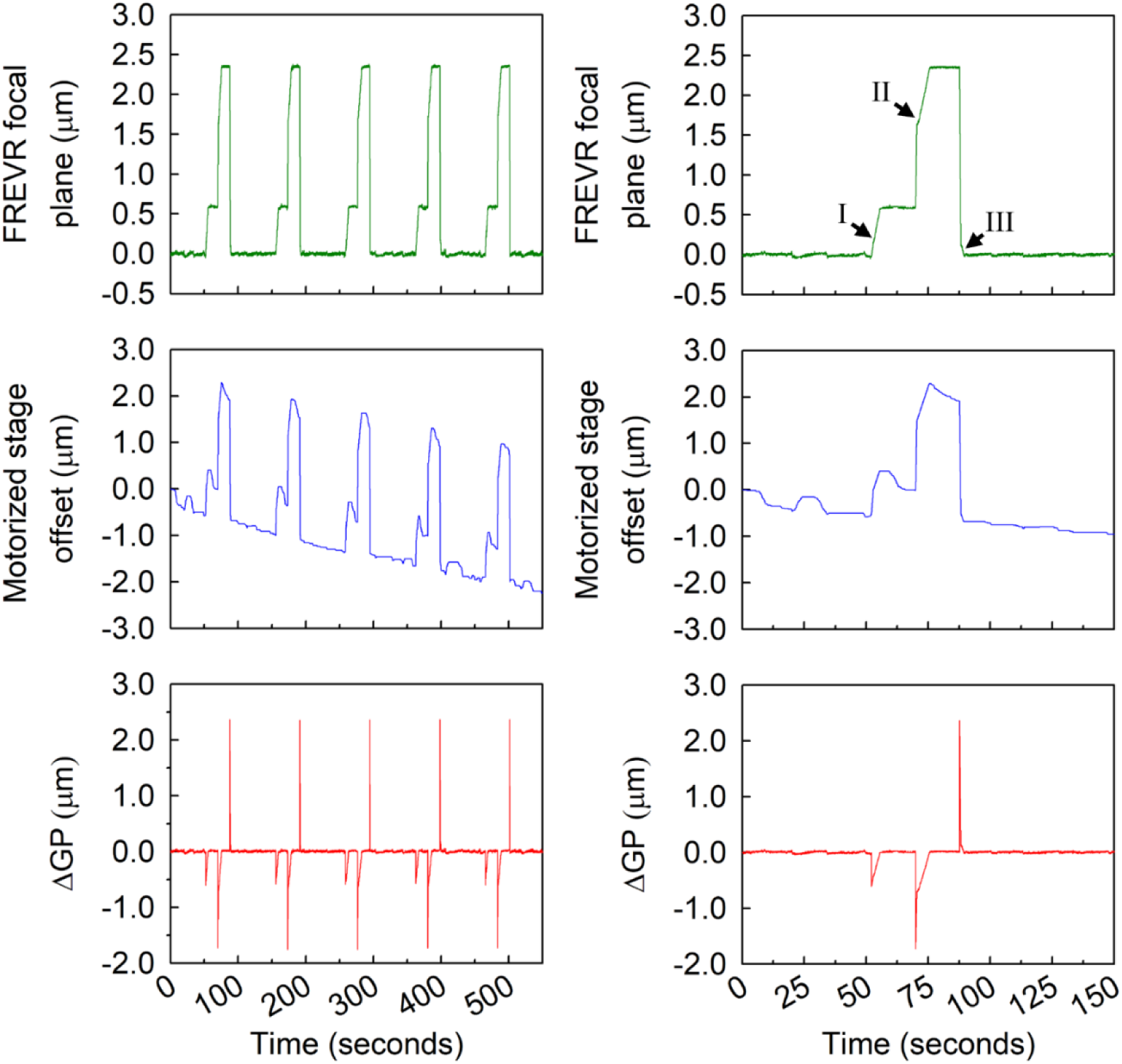
Traces obtained from FREVR show fiducial bead position (top), motorized stage offset readout from microscope (middle), and deviation between desired and measured focal planes (ΔGP, bottom) over five sequential z-scans. Left panels show the full experiment and right panels show a zoomed view of the first z-scan, with markers indicating stage repositioning events and uncorrected axial drift (further details in Figure S3).

It should be emphasized that the FREVR-assisted positioning method described here provides highly precise and repeatable alternation between imaging planes. While the accuracy of the reported axial positions is limited by the determination of the axial calibration factor, it is nevertheless inconsequential for the present study, which relies on precise repeated sampling of the same planes rather than their nominal axial coordinates.

#### 2.4.3 Cell samples imaged with FREVR

Fields containing cells co-expressing mEGFP-M_2_R and mCitrine-Arr2 were identified, and a nearby RB, attached to the dish surface, was selected for FREVR calibration. Prior to imaging, a reference library for the RB was generated from a bead imaging sequence spanning 4.68 µm in steps of 23 nm. An exploratory FREVR-driven z-stack of fluorescence imaging was performed in the vicinity of the basolateral membrane to precisely determine the focal plane of the basolateral membrane of the cells in the imaging field. The plane corresponding to the basolateral membrane, which was determined by finding the maximum intensity within the exploratory FREVR-driven z-stack, was set as the starting plane for FREVR-enabled z-scans.

Each z-scan contained a scan performed at the basolateral membrane, and one +2.34 µm above the basolateral membrane. Z-scans were performed in five-minute intervals to reduce the effects of photobleaching. After two untreated z-scans, the M_2_R agonist carbachol was added to a final concentration of 300 µM shortly before the third z-scan. Subsequent z-scans were performed in five-minute intervals for one hour after the addition of the ligand.

### 2.5 Unmixing of spectrally resolved fluorescence images

Spectrally resolved fluorescence images containing pixel-level composite emission spectra comprised of signal from both mEGFP and mCitrine were unmixed using the custom-built software package OptiMiS DC^29^. Elementary emission spectra of mEGFP and mCitrine were determined from cells expressing only mEGFP-M_2_R or mCitrine-Arr2 respectively. These elementary emission spectra of mEGFP and mCitrine were used to deconvolute (unmix) the composite spectra contained within each pixel; enabling pixel-level signal-quantification of each protein, as described previously^30–32^. The unmixed data were saved into separate three-dimensional fluorescence image stacks, two spatial dimensions and one temporal dimension, for downstream spatial-temporal analysis.

### 2.6 Basolateral image visualization and colocalization mapping

Basolateral membrane image stacks were normalized to facilitate visual comparison of receptor and arrestin distributions across time and between cells (Figure 3). For each image stack, pixel intensities were normalized by dividing all pixel values by the intensity of a representative high-signal punctum exhibiting clear colocalization of mEGFP-M_2_R and mCitrine-Arr2.

**Figure 3.**
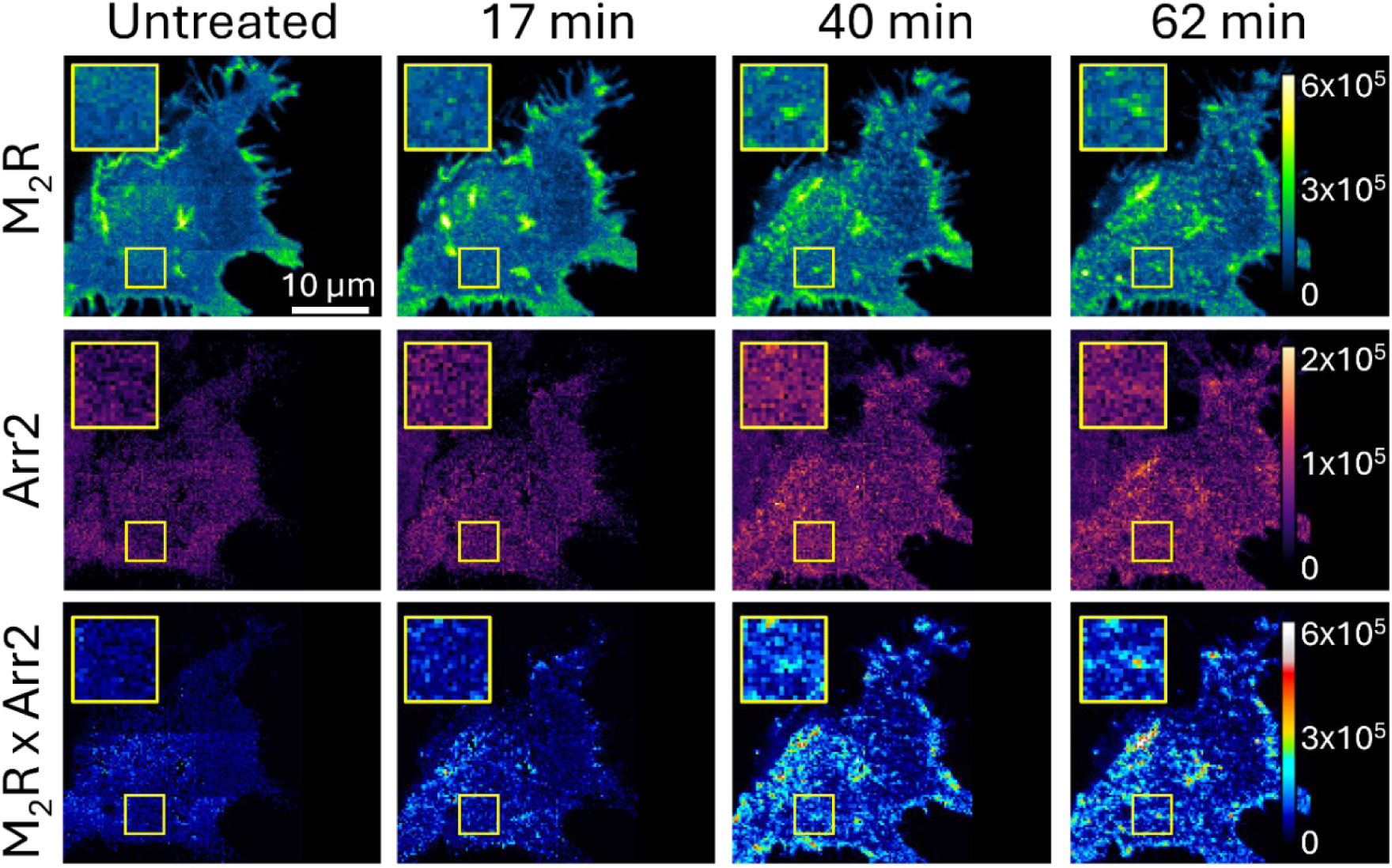
Typical fluorescence micrograph time series at the basolateral membrane of a single cell co-expressing mEGFP-M_2_R and mCitrine-Arr2 before and after agonist stimulation. Measured fluorescence intensity of mEGFP-M_2_R (top), mCitrine-Arr2 (middle), and composite (bottom) reveal changes in the local protein environment which are most apparent in regions where the membrane appears uniform prior to stimulation but becomes punctated following ligand treatment, as illustrated within the yellow demarcated region of interest. Fluorescence intensity values were assigned false colors according to scales shown on the right side of each row.

Following normalization, pixel-wise multiplication of the mEGFP-M_2_R and mCitrine-Arr2 image stacks was performed to generate qualitative colocalization maps highlighting regions of concurrent receptor and arrestin localization. The normalized M_2_R, Arr2, and colocalization images were visualized using fixed lookup tables applied consistently across datasets: blue-green-yellow for M_2_R, purple-orange-yellow for Arr2, and blue-red-white for the multiplied colocalization maps. These visualization steps were used solely for qualitative assessment and presentation and were not used for quantitative analysis.

### 2.7 Quantitative modeling of cross-section fluorescence profiles

Cross-section images were inspected to identify regions with smooth membrane cross-section and uniform cytoplasmic mCitrine-Arr2 fluorescence. A region of interest (ROI) was drawn to span from the cytoplasm, across the membrane, and into extracellular space (Figure 4). For each ROI, the M_2_R images were analyzed by identifying, in each row, the column position of the maximum fluorescence intensity corresponding to the plasma membrane. The rows were then shifted to align all peak-intensity pixels into a single column, producing a matrix in which intracellular, membrane, and extracellular regions were spatially aligned. This alignment reduced curvature effects and enabled averaging over a larger portion of the cell for improved statistics. The same organization matrix determined from the M_2_R dataset was applied to the Arr2 images to preserve spatial correspondence. Mean fluorescence intensity was then computed for each column, yielding one-dimensional fluorescence intensity profiles as a function of pixel position.

**Figure 4.**
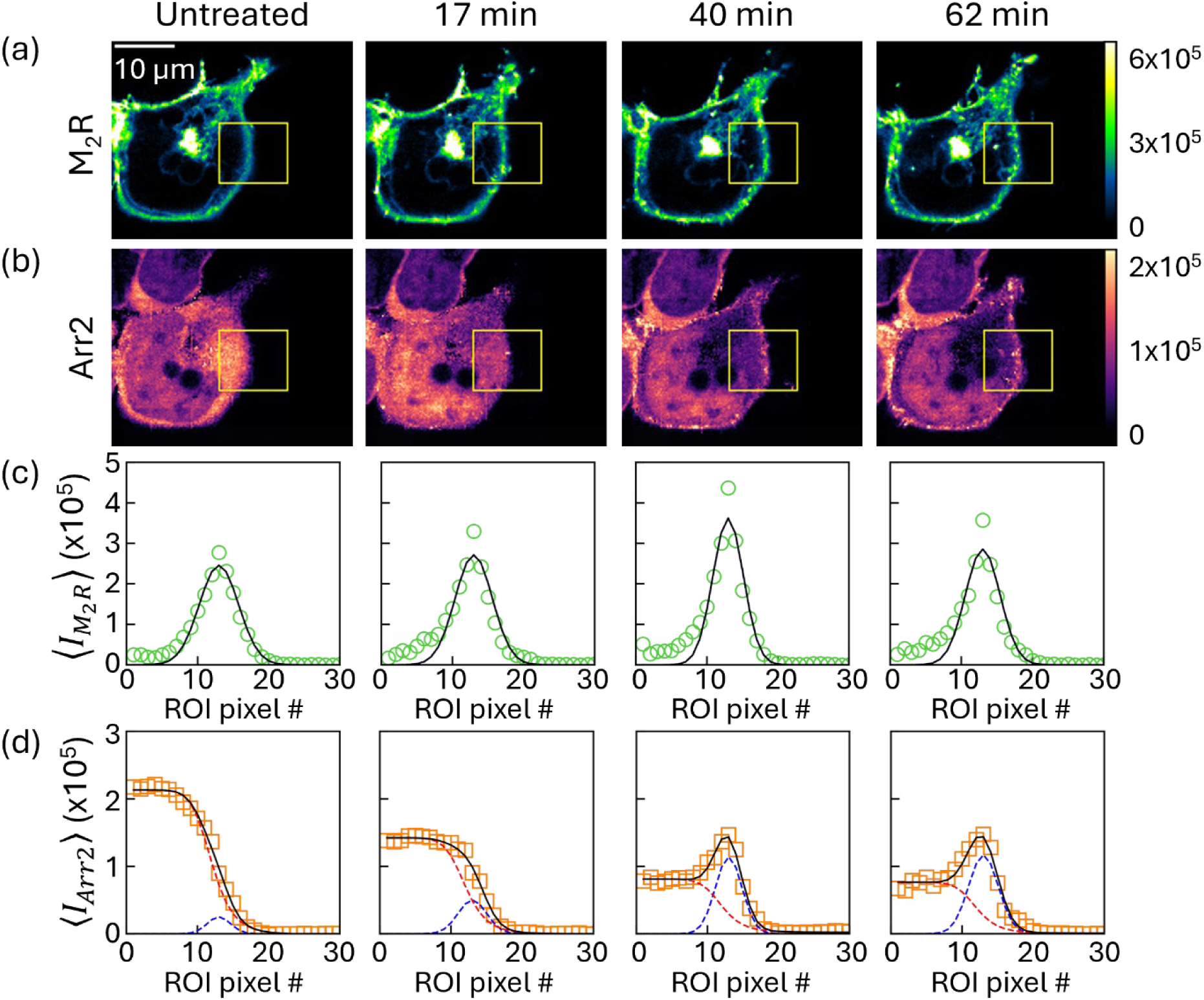
(a,b) Fluorescence micrographs of cellular cross-sections showing mEGFP-M_2_R (top) and mCitrine-Arr2 localization (bottom) before ligand treatment and at selected time points after activation; M_2_R remains primarily membrane-localized, while Arr2 redistributes from the cytoplasm to the plasma membrane. Yellow square represents the ROI used for further analysis. (c,d) Corresponding curvature-corrected intensity profiles for mEGFP-M_2_R (top) and mCitrine-Arr2 (bottom) extracted from the region of interest in (a,b), illustrating increased-membrane-associated Arr2 following receptor activation. The black lines represent fits using eq 1 in (c) and eq 1 + eq 2 in (d) as detailed in Methods Section 2.7.

Each fluorescence intensity profile was modeled to quantify membrane-associated and cytoplasmic components. Profiles were fit with either a Gaussian function (in the case of the M_2_R image stack) or a combination of Gaussian and sigmoidal functions (in the case of Arr2 image stack). The membrane component was fit with a Gaussian function defined as:

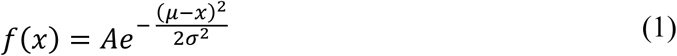

Where *x* is column position, A is the amplitude of the intensity profile, µ is the aligned membrane column position (held fixed during fitting), and σ the width of the Gaussian. Cytoplasmic fluorescence was described using a sigmoidal function:

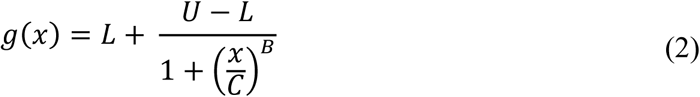

where L and U represent the lower and upper limits of the intensity profile, respectively, B represents the slope of the intensity profile from the upper limit to the lower limit, and C represents the column position of the inflection point.

Profile fitting was performed using custom Python software which applied a nonlinear least-squares optimization algorithm until convergence. The curvature-corrected mEGFP-M_2_R maps were fitted using eq 1, with µ fixed to the column corresponding to the aligned membrane position, while *A* and σ were allowed to vary during fitting. When fitting the curvature-corrected mCitrine-Arr2 maps, the sum of eq 1 and eq 2 was used. As with the mEGFP-M_2_R maps, µ was fixed according to the aligned membrane column. The parameter C of the sigmoidal function, corresponding to the position of the inflection point, was first determined from fits to untreated images of a particular time series. The resulting C values from the untreated images in a given time series were averaged and subsequently fixed when refitting the remaining datasets to maintain consistency across experimental conditions.

The resulting fits were used to extract quantitative parameters describing membrane-localized fluorescence (Figure 5). The signal emitted from the membrane was determined by integrating the Gaussian function, given by:

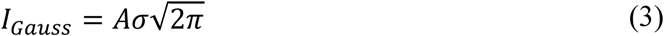

**Figure 5.**
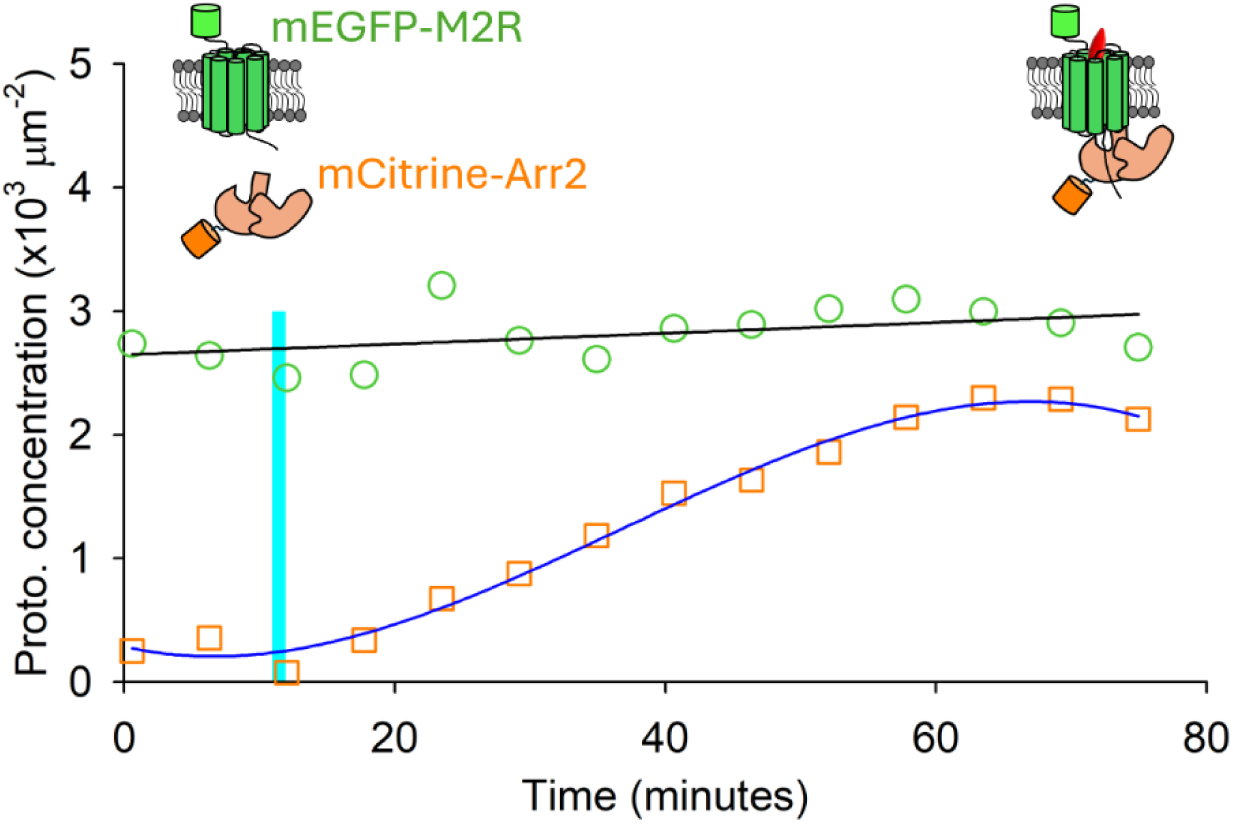
Concentration of membrane-associated M_2_R (green circles) and Arr2 (orange squares) plotted against time. M_2_R and Arr2 data were fitted with a linear regression (black line) and a third-order polynomial (blue line), respectively, to capture the overall trends and serve as guides to the eye. Cartoon representations illustrate the system before and after ligand addition: mEGFP-M_2_R (green GPCR with attached fluorophore), mCitrine-arrestin-2 (orange protein with attached fluorophore), and ligand (red oval).

This value was then normalized by the fluorophore brightness (ε^proto^) and an area factor (∭ PSF^2^(0, *y*, *z*)*dydz*) to yield an effective concentration of membrane-associated molecules in the following form:

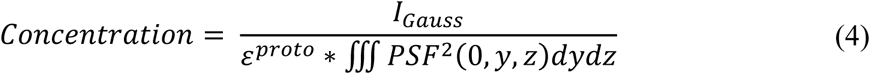

where the integral of the PSF^2^ was computed numerically as described previously^28^.

### 2.8 Determination of monomeric brightness

To determine monomeric brightness (ε^proto^) of each fluorescent protein, purified mEGFP and mCitrine solutions were diluted in HEPES buffer with 1 mM DTT and imaged using the same fluorescence microscope described in Methods Section 2.3. Solutions of mEGFP and mCitrine were measured at multiple concentrations (7.7, 3.8, 1.9, 1.0, 0.5 µM). Fluorescence intensity data were plotted against respective molecular concentrations and fit by linear regression. The slope of the linear regression, Θ, was used to calculate ε^proto^ following eq S22 provided in the Supporting Information of Stoneman et al.^28^:

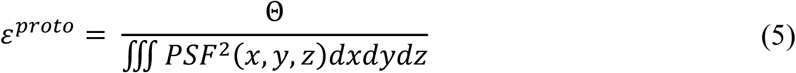

where the integral of the PSF^2^, representing the volume within the excitation beam, was determined numerically, similarly to the previous section.

## 3. RESULTS

### 3.1 Instrument Design and Experiments

To monitor the localization and kinetics of M_2_R and Arr2 at the cellular level, we focused on two specific regions of the plasma membrane: the basolateral membrane, which affords a frontal view of the receptor oligomerization and clustering into dense puncta (most likely, clathrin-coated pits), and a second plane located 2.34 μm above the basolateral membrane, where we can better visualize arrestin’s initial localization in the cytoplasm and its move towards the membrane following receptor activation or away from it during its eventual re-internalization. To track dynamic changes over time, we repeatedly alternated between imaging these two planes throughout the experiment. This interleaved scanning approach allowed us to monitor events at the membrane and in the adjacent cytoplasm in near real-time, capturing both receptor clustering and arrestin translocation with sufficient time resolution. While the basolateral view provides detailed insight into membrane-bound processes, including receptor localization, interactions, and the formation of puncta, the cellular cross-section provides a better view of cytoplasmic proteins like arrestins and their association with the plasma membrane. Achieving reliable alternation between these imaging planes requires precise and repeatable axial positioning.

The FREVR system, detailed in Methods Section 2.3, was implemented in our instrument using fiducial reference beads for real-time imaging. In our setup, fiducial reference beads (RB), a ∼3.75 µm diameter amino-coated polystyrene bead, were attached to the sample coverslip for focal position tracking (Figure 1a). The RBs were illuminated by a 625 nm light-emitting diode (LED), and the reflected bead signal was detected by a Complementary-Metal-Oxide-Semiconductor (CMOS) camera. Before imaging, a reference library, or image stack, was collected for a selected bead by capturing a series of images at 23 nm axial intervals spanning the focal range of interest (4 microns), as detailed in Methods Section 2.4. This high-resolution stack served as a lookup table, enabling the system to determine the bead’s current focal plane by comparing each live image to the pre-recorded reference frames. The resulting positional information was used to control the motorized stage of a Zeiss Axio Observer microscope.

To implement FREVR on our fluorescence microscope we adapted the optical layout to suit our instrument. We introduced a new optical pathway by adding a notch dichroic mirror (DM) that reflects 633 nm light while allowing both longer and shorter wavelengths to pass through. This enabled us to create a dedicated detection channel for bead tracking, separate from the fluorescence imaging path. Together, these modifications enabled real-time monitoring and correction of focal drift, resulting in a fully functional FREVR implementation shown in Figure 1b.

### 3.2 Mitigation of Focal Plane Positioning Errors and Temporal Drift Using FREVR

To quantify the axial positioning limitations of our system, we performed a series of controlled z-positioning tests using the FREVR-equipped microscope described in Section 3.1 and sample chambers containing fiducial reference beads. In a representative experiment, five sequential z-scans were performed over a ten-minute period. Each z-scan consisted of three imaging planes: a baseline plane, a secondary plane positioned 0.59 µm above the baseline, and a third plane positioned 2.34 µm above the baseline. During the experiment, we simultaneously recorded three data streams (Figure 2): the measured focal plane position determined by FREVR, the motorized stage offset reported by the Zeiss Axio Observer, and the difference between the measured focal position and the target position commanded by the control program at any given instant. When the objective is being moved between planes, this measured–target difference primarily reflects dynamic effects (lag, settling, overshoot) as the objective moves to the new setpoint; during stationary holds, it reflects true focal drift.

We observed that the microscope’s motorized stage consistently undershot the desired position when commanded to move to the next step in the z-scan, likely due to mechanical backlash in the translation stage drive (Figure 2). FREVR detected and actively corrected these discrepancies in real time, as highlighted at positions marked with I, II, and III in Figure 2. Moreover, across the full ten-minute scan series, the motorized stage trace further revealed that continuous vertical adjustments were required to maintain alignment with the target focal planes. Without compensation, the cumulative drift during this time would have reached ∼2 µm; however, with FREVR engaged, these displacements were effectively mitigated. Across all measurements, the intrinsic error in axial positioning while FREVR was engaged remained within tens of nanometers, demonstrating highly stable and precise focal control over extended imaging sessions.

These benchmark results validated FREVR as a critical component for accurate live-cell imaging, especially when imaging multiple planes repeatedly over time. With this stability in place, we next applied this imaging setup to HEK-293 cells expressing fluorescently tagged Arr2 and M_2_R. A scanning protocol was designed to monitor two planes of interest, the basolateral membrane and cross-section 2.34 µm deep into the cell, before and after activation of M_2_R by an agonist ligand. In the sections that follow, we first present results from the basolateral membrane, followed by a detailed analysis of intracellular dynamics observed in cellular cross-sections.

### 3.3 Visualization of M_2_R and Arr2 Dynamics at the Basolateral Plasma Membrane Following Receptor Activation

Spectrally resolved fluorescence images of cells expressing mEGFP-M_2_R and mCitrine-Arr2 were obtained using the two-photon optical micro-spectroscope described in detail in Methods Section 2.3. This system captures full emission spectra at each pixel, rather than filtering fluorescence into fixed channels, enabling more precise resolution of overlapping fluorescent signals. To separate the contributions from each fluorophore, we applied a previously described spectral unmixing algorithm that operates at the pixel level^30–32^, which is also described succinctly in Methods Section 2.5. Unmixing of the composite images resulted in two distinct spatial maps: one representing the fluorescence intensity of mEGFP-M_2_R, and the other representing mCitrine-Arr2. This allowed us to track the distribution of each protein independently within the same cell over time.

To monitor dynamic changes in protein localization, we acquired a time series of fluorescence images consisting of fourteen consecutive z-scans, each separated by a five-minute interval. This time interval was selected to balance the need for capturing time sensitive cellular changes while reducing the effects of photobleaching. In each z-scan, fluorescence was recorded at two predefined axial positions: the basolateral membrane and a cross-section 2.34 µm above it. Samples were imaged prior to and after the addition of the M_2_R agonist carbachol^33^ to examine how receptor activation influences M_2_R membrane organization and its interactions with Arr2.

Fluorescence scans performed at the basolateral membrane revealed pronounced changes following receptor activation, as shown by the representative sample in Figure 3. Prior to addition of the agonist ligand, the mEGFP-M_2_R fluorescence appeared bright and mostly uniformly distributed across the membrane, aside from a few larger artifacts which appear consistent with membrane inhomogeneities (folds, invaginations, etc.). In contrast, the mCitrine-Arr2 maps display low levels of fluorescence signal, consistent with a primarily cytoplasmic distribution (i.e., localized above the membrane patch being imaged) and minimal membrane association. Upon agonist stimulation, mCitrine-Arr2 fluorescence increased noticeably in overall intensity, indicating Arr2 recruitment to the membrane. In both mEGFP-M_2_R and mCitrine-Arr2 fluorescence intensity maps, we observed the emergence of discrete puncta, indicative of the formation of punctate membrane structures consistent with clathrin-coated pits (CCPs) and other endocytic structures^34^. Time-lapse imaging showed that many of these assemblies formed, migrated, and dissipated over minutes after treatment. To highlight regions where M_2_R and Arr2 co-localize, we calculated a pixel-wise product of the two fluorescence maps, generating a composite image enhancing visibility of colocalization between the two proteins of interest, as detailed in Methods Section 2.6.

These results indicate significant biological activity occurring after ligand stimulation, including the colocalization of M_2_R and Arr2 and the formation of CCPs and other endocytic structures. However, imaging only the basolateral membrane limits interpretation, as the observed increase in Arr2 fluorescence could arise either from enhanced membrane association or from an overall increase in cytoplasmic Arr2 levels. To resolve this ambiguity and distinguish between these possibilities, we next examined cellular cross-sections to assess changes in cytoplasmic and membrane-bound Arr2.

### 3.4 Visualization of M_2_R and Arr2 Spatial Redistribution After Activation of M_2_R as Observed in Cellular Cross-Sections

All 2.34 µm deep cellular cross-section images obtained from the time-lapse z-scans described in the previous section were spectrally unmixed to generate separate fluorescence intensity maps for mEGFP-M_2_R and mCitrine-Arr2. Visual analysis of the resulting time-resolved 2D maps revealed that M_2_R was primarily localized to the plasma membrane both before and after ligand treatment. In contrast, Arr2 displayed dynamic redistribution, shifting from a predominantly cytoplasmic pattern in untreated cells to a more membrane-associated localization following M_2_R activation.

To quantify changes in protein localization, a region of interest (ROI) was defined in each cross-sectional image to span from the cytoplasm, across the plasma membrane, and into the extracellular space outside the cell, as shown by the yellow rectangle in Figure 4. The fluorescence data within the ROI were then spatially reorganized according to the protocol described in Methods Section 2.7, aligning the membrane positions into a common reference column generating a curvature-corrected data matrix. Rows containing obvious vesicles or endocytic structures, which could skew the determination of the plasma membrane peak, were excluded from the curvature alignment step to ensure accurate detection of the underlying plasma membrane position. After alignment, fluorescence intensities were averaged by column within the ROI, producing a curvature-corrected intensity profile for each protein at each time point.

The resulting curvature-corrected intensity profiles were analyzed using mathematical models selected to reflect the distinct spatial distributions of M_2_R and Arr2 (see Methods Section 2.7 for full details). The fluorescence profiles of mEGFP-M_2_R were modeled using a membrane-localized component only, due to the strong membrane-localization behavior of the receptor. In contrast, mCitrine-Arr2 profiles required a composite model consisting of a membrane-localized component and a cytoplasmic contribution.

Figure 4 illustrates the modeled intensity profiles of both proteins of interest, revealing minor intensity fluctuations of mEGFP-M_2_R at select treatment times while the mCitrine-Arr2 fitted curves show a pronounced decrease in cytoplasmic Arr2 and a corresponding increase in membrane-associated Arr2 following receptor activation. These modeled intensity profiles support the visual analysis, confirming the redistribution of Arr2 toward the plasma membrane and motivating further quantification of protein concentrations at the membrane.

Membrane-associated protein concentrations were calculated using eq 4 described in Methods Section 2.7. The estimated membrane-associated protein concentrations were plotted against treatment time (Figure 5). M_2_R concentration remained largely constant throughout the experiment, whereas membrane-associated Arr2 increased markedly following agonist treatment, consistent with the visual observations described earlier. This increase in membrane-associated Arr2 is consistent with the recruitment of cytoplasmic arrestin to activated receptors at the plasma membrane, as expected for GPCR-arrestin interactions. The relatively stable M_2_R concentration suggests that receptor redistribution does not substantially alter the total membrane-associated receptor population on the timescale of these measurements. These results demonstrate that membrane-associated protein concentrations can be quantitatively monitored in living cells over time, providing a foundation for more detailed analyses of receptor-arrestin interaction dynamics.

## 4. CONCLUSIONS

The implementation of FREVR on our two-photon microscope equipped with a standard motorized objective stage revealed substantial improvements in both the accuracy and precision of axial sample plane positioning during live-cell imaging over extended periods of time. In this implementation, image-forming sample planes spanning approximately 3 µm could be revisited with a precision on the order of ∼20 nm, enabling reproducible sampling of multiple axial positions. Although demonstrated here using a two-plane imaging strategy targeting the basolateral membrane and a cellular cross-section, the same principle is expected to improve the reliability of full z-stack acquisitions used for high-resolution three-dimensional reconstructions in any laser-scanning microscope. In this regard, FREVR ensures that successive focal z-stack steps correspond to well-defined and reproducible spatial positions.

Using this spatially stabilized imaging approach, we monitored the dynamic redistribution of the muscarinic acetylcholine M_2_ receptor and arrestin-2 in living cells following receptor activation. Imaging at the basolateral membrane revealed spatial reorganization of receptors into punctate regions accompanied by increased colocalization of arrestin. At the same time, cross-sectional imaging showed that initially cytoplasm-distributed arrestin progressively accumulated at the plasma membrane following stimulation, consistent with arrestin migration and binding to activated receptors. Unlike conventional approaches that rely on measurements from separate cells or independently acquired focal planes, this method enables direct, time-resolved comparison of membrane and cytoplasmic dynamics within the same cell. By minimizing the variability arising from cell-to-cell heterogeneity and focal drift, this approach provides a more reliable framework for quantifying receptor-arrestin interactions in living cells. The ability to repeatedly access defined imaging planes with nanometer-scale precision should facilitate further investigations of receptor-arrestin dynamics and related signaling processes. In particular, improved axial stability will enable more detailed quantitative analyses of the kinetics, spatial organization, and stoichiometry of signaling complexes at the plasma membrane. More broadly, approaches that enhance positional accuracy during fluorescence imaging will help support increasingly quantitative investigations of protein localization and interactions in living cells.

## ASSOCIATED CONTENT

### Data Availability Statement

Spectrally resolved fluorescence images were unmixed using an in-house software called OptiMiS-DC^29^, accessible via a community repository (https://doi.org/10.5281/zenodo.15588222).

Further information or data are available from the authors upon reasonable request.

### Supporting Information

The Supporting information is available free of charge at:

Detailed derivation and experimental validation of axial stage calibration using fiducial beads, including determination of the axial correction factor. Graphical illustration of vertical plane correction and stabilization performed by FREVR in a representative experiment containing multiple sampled planes.

## AUTHOR INFORMATION

### Notes

The authors declare no competing financial interest.

## ACKNOWLEDGEMENTS

The authors thank undergraduate students, Rebecca J. June and Melissa Gleiter, from the University of Wisconsin-Milwaukee (UWM) for their contributions to cell cultures and data analysis.

## ABBREVIATIONS

HEK-293: human embryonic kidney 293
GPCR: G protein-coupled receptor
Arr2: arrestin-2
M_2_R: muscarinic acetylcholine M_2_ receptor
FREVR: focal readjustment for enhanced vertical resolution.

