## SupplementaryMaterials for "Precise Alternation Between Image-Forming Sample Planes Enables Quantitative Monitoring of Receptor-Arrestin Interaction Dynamics at the Plasma Membrane of Live Cells"

**Supplementary Note 1**

To calibrate the objective position readouts from the Zeiss Axio Observer, we performed experiments using two different types of polystyrene beads of known size glued to glass coverslips (See Section 2.4.1). In these experiments, maximum values of the bead radii were determined from horizontal and vertical reflected-light intensity profiles extracted from reflected-light images acquired at successive objective positions. The reported objective position at which the intensity profile reached its maximum width (in pixels) was taken tentatively to correspond to the focal plane passing through the bead center; this assumption will be tested below. Two measurement strategies were employed: (A) determining the difference in reported objective position from the glass surface to the bead center, and (B) determining the reported axial position between the centers of two beads of different sizes. To account for any discrepancies between measured objective-position differences and true focal plane displacements, we introduced an axial correction factor, $\alpha$ (See Section 2.4.1). This factor relates bead radii estimated from reported objective positions to their known physical radii. For each bead, we define:

$$\alpha_1 r_1 = R_1, \tag{1}$$

$$\alpha_2 r_2 = R_2, \tag{2}$$

where $\alpha_1$ and $\alpha_2$ are correction factors, $r_1$ and $r_2$ are the experimentally measured radii, and $R_1$ and $R_2$ are the manufacturer-reported bead radii (Spherotech and Dynabeads). Separate correction factors were initially allowed for each bead to account for potential size- or shape-dependent effects. To estimate these correction factors, we employed two independent methods based on 3D calibration stack measurements (Section 2.4.1, Figure S1, and S2).

Method A

The bead radius was determined by measuring the objective-reported distance between the glass surface and the bead center. The bead center was identified as the focal plane corresponding to the maximum bead diameter in cross-section images, while the glass surface was identified as the plane corresponding to the maximum amplitude of a surface impurity (Figure S1). Solving eq 1 and 2 for $\alpha_n$ gives:

$$\alpha_n^A = \frac{R_n}{r_n}. \qquad (3)$$

where the index letter *A* denotes the correction factor was determined using Method A. Using $R_n$ values from manufacturers' specifications and $r_n$ values averaged from five individual experiments, we obtained $\alpha_1^A = 1.17 \pm 0.15$ from Spherotech beads and $\alpha_2^A = 1.17 \pm 0.18$ from Dynabeads.

Method B

An independent estimate of the calibration factor was obtained by measuring the objective-reported separation between the centers of two beads of different sizes (Figure S2). The correction factor from these experiments can be calculated by dividing the difference in known bead radii by the difference in the measured bead radii:

$$\alpha^B = \frac{R_1 - R_2}{r_1 - r_2}. \qquad (4)$$

This correction factor, $\alpha^B$, was determined experimentally by averaging data from five independent experiments, yielding $\alpha^B = 1.16 \pm 0.54$, which agrees with the axial correction factors obtained from Method A.

To rigorously show that the correction factors should not be bead dependent, we can use the fact that $\alpha^B = \alpha_1^A$ and we can substitute $\alpha_1^A$ into eq 4.

$$\alpha_1^A = \frac{R_1 - R_2}{r_1 - r_2}. \tag{5}$$

Multiplying both sides by $r_1 - r_2$ gives:

$$\alpha_1^A r_1 - \alpha_1^A r_2 = R_1 - R_2. \tag{6}$$

Which can be rewritten by substituting in equations (1) and (2) on the right side of the equation:

$$\alpha_1^A r_1 - \alpha_1^A r_2 = \alpha_1^A r_1 - \alpha_2^A r_2. \tag{7}$$

By subtracting $\alpha_1 r_1$ from both sides, we find:

$$-\alpha_1^A r_2 = -\alpha_2^A r_2. \tag{8}$$

By dividing both sides by $-r_2$ it is revealed that

$$\alpha_1^A = \alpha_2^A. \tag{9}$$

This identity was true experimentally, but here we show that it should retain its validity with the assumptions made in this derivation.

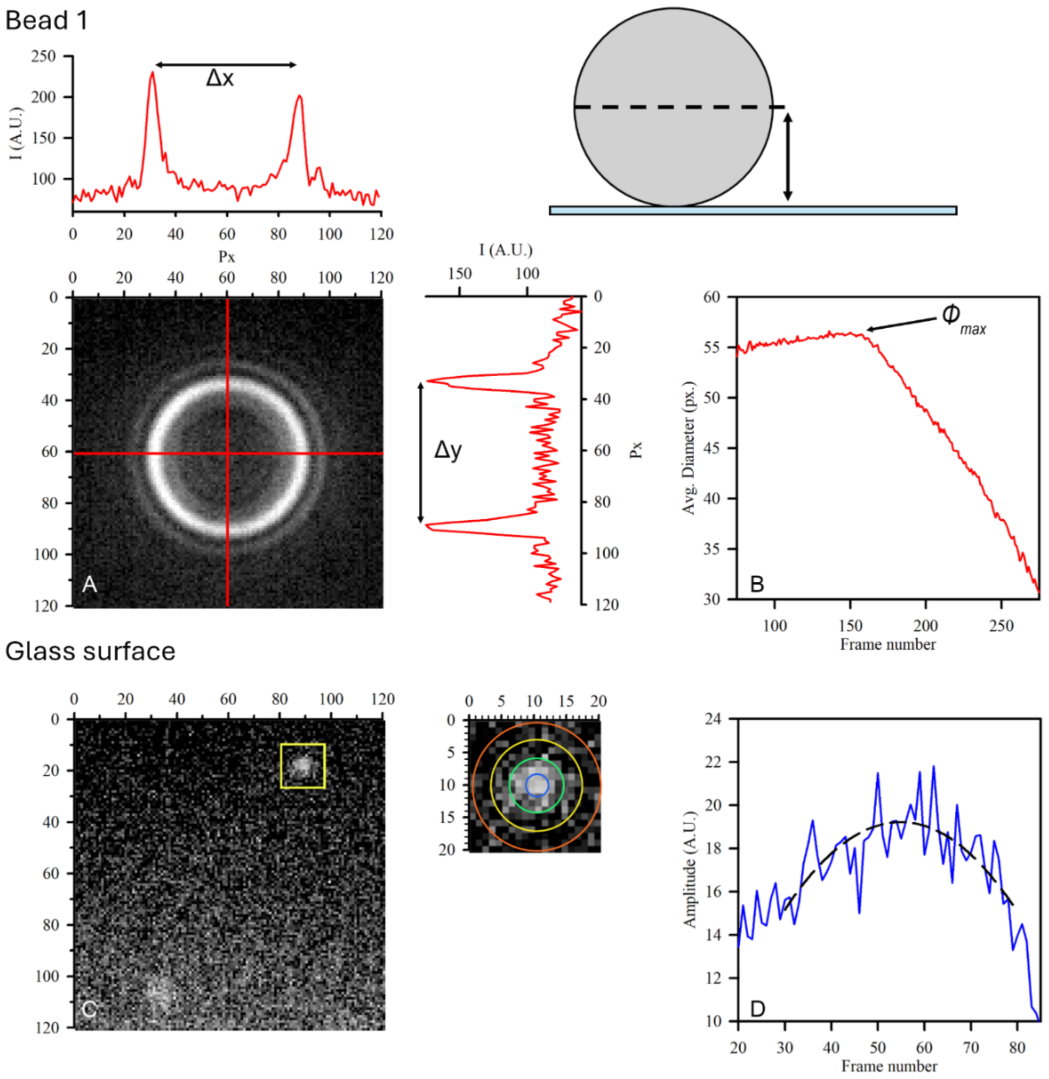


**Figure S1**. Microscope Z stage calibration using distance from glass surface to center of bead (Method A: absolute radius measurement). (A) Single frame of a 3D stack of reflected bead images of a single bead. The corresponding horizontal and vertical intensity profiles extracted through this bead are plotted to the top and to the right of the reflected bead image, respectively (A). The horizontal (Δx) and vertical (Δy) widths of the bead center, in pixels, were obtained from double-Gaussian fits of each of the intensity profiles. The average diameter of the bead center, $\phi_j$, was computed as the mean of Δx and Δy for each frame $j$ in the 3D image stack. The average diameter, $\phi_j$, was plotted as a function of frame number to determine the frame number of maximal diameter ($\phi_{max}$) corresponding to the bead center (B). A representative glass surface image (C) shows a region of interest (yellow box) containing a surface impurity. This region was fitted with a two-dimensional Gaussian to identify the focal plane of maximum intensity, corresponding to the glass surface (D). The reported objective distance between the frame number of the glass surface and the frame number corresponding to $\phi_{max}$ was used to determine the bead radius, $r_1$ , which was compared to the known bead radius, $R_1$ , to compute the axial correction factor (Supplementary Note 1, Method A).

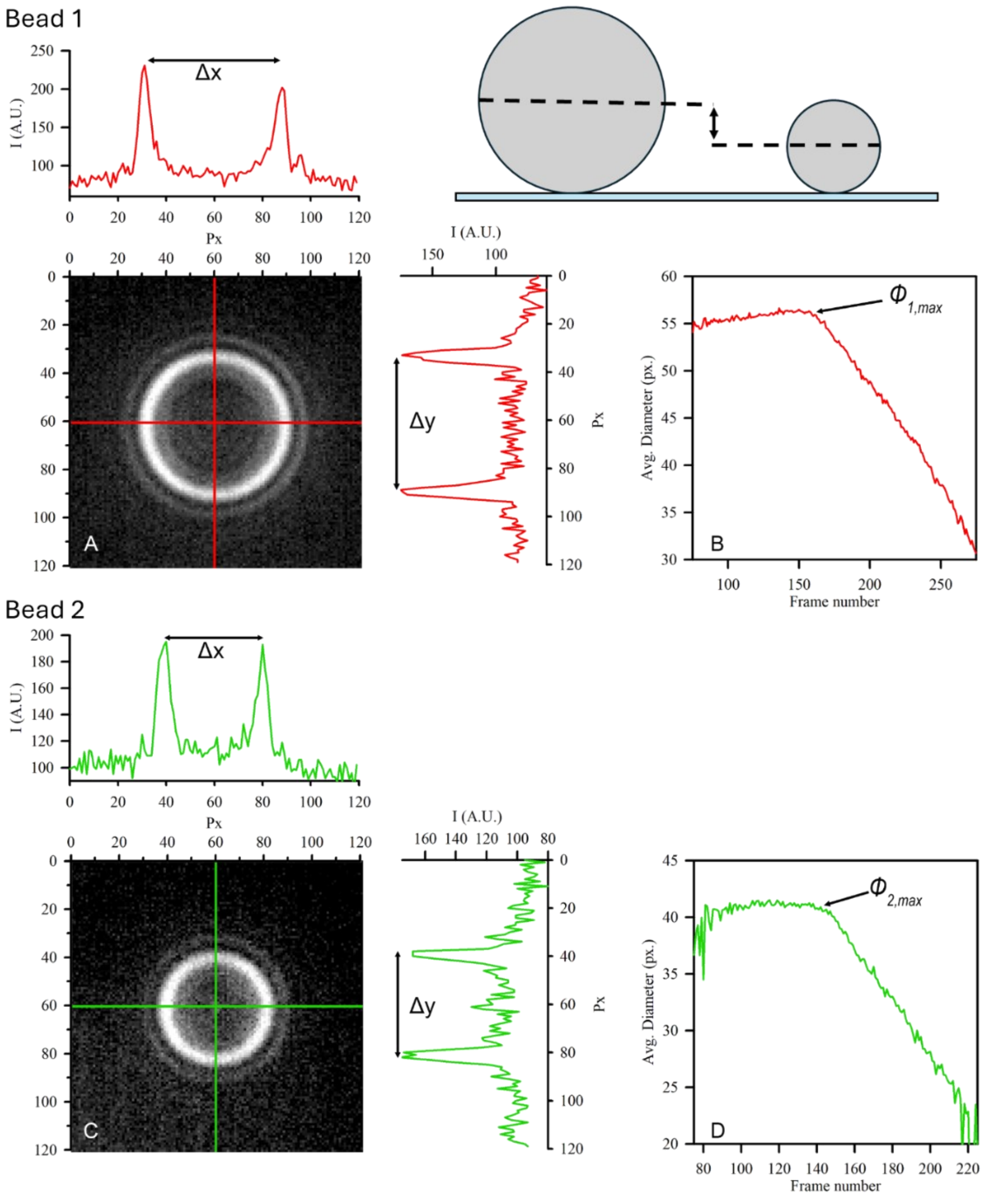


**Figure S2**. Microscope Z stage calibration using distance between the center of two different sized beads (Method B: relative radius measurement). Representative images of bead 1 (A) and bead 2 (C) with corresponding intensity profiles above and to the right of each bead image (A, C). Bead diameters (Δx, Δy) were determined from double-Gaussian fits, and the average bead diameter was calculated as the mean of Δx and Δy. The focal plane corresponding to the maximum diameter was identified for each bead ($\phi_{1,max}$, B; and $\phi_{2,max}$, D). The reported objective distance between these focal planes was used to estimate the difference in bead radii. This measured difference was compared to the known difference in bead radii to compute the axial correction factor (Supplementary Note 1, Method B).

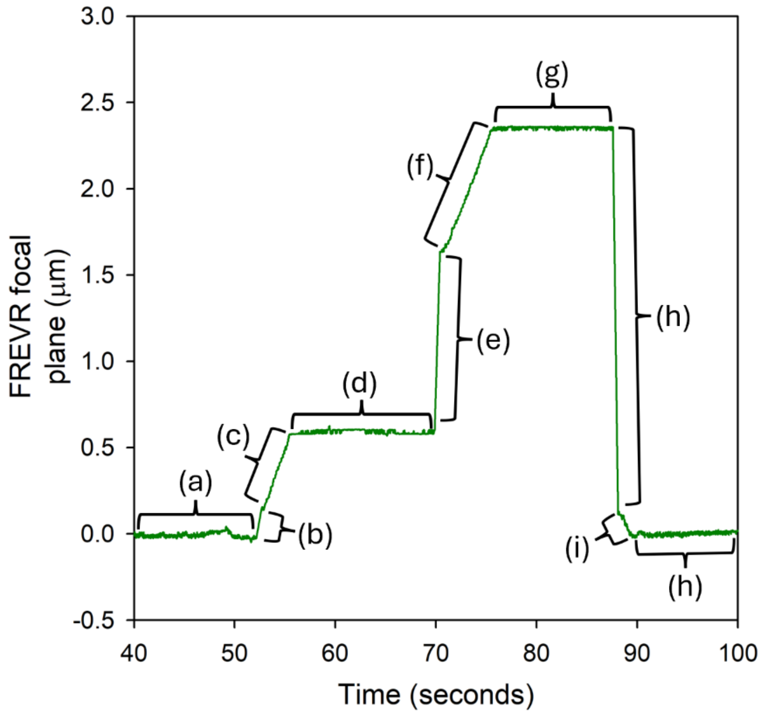


**Figure S3**. A representative trace demonstrates FREVR stabilization and compensation for mechanical and optical instabilities. (a) FREVR maintains the desired focal plane until a command is given to move. (b) The microscope stage is given a command to move +585 nm but undershoots its target due to microscope backlash. (c) FREVR actively corrects the position until the appropriate focal plane is reached. (d) The position is maintained until the next command. (e) The command for a second step of +1,755 nm is given, and (f) FREVR actively corrects to the appropriate position. (g) The plane is maintained until (h) the microscope is given the command to return to the original position, and (i) only a slight correction is needed by FREVR until it reaches (h) the desired plane.
